# Cardiac defects in spinal muscular atrophy and the role of SMN in cardiomyocyte homeostasis

**DOI:** 10.64898/2026.03.20.713246

**Authors:** Reid Garner, Leillani L. Ha, Flavia C. Nery, Rebecca G. Spellman, Lucia Chehade, Benjamin Glennon, Eric J. Eichelberger, Salomé Da Silva Duarte Lepez, Alec J. Johnstone, Rashmi Kothary, Kathryn J. Swoboda, Christiano R. R. Alves

## Abstract

Spinal muscular atrophy (SMA) is characterized by motor neuron degeneration caused by deficiency of the survival motor neuron (SMN) protein. However, evidence increasingly supports broader systemic involvement. This study aimed to examine cardiac pathology in SMA patients and to investigate how reduced SMN levels impact cardiomyocyte homeostasis. We analyzed postmortem data from 14 SMA type I patients from the pre-treatment era, integrating gross anatomical, histopathological, and clinical findings. To investigate cardiomyocyte-intrinsic effects of SMN deficiency, healthy human cardiomyocytes were subjected to SMN knockdown and assessed using metabolic assays and transcriptomic profiling. Key findings were further investigated in vivo using the *Smn^2B/−^* mouse model of SMA. We found heterogeneous cardiac involvement in SMA patients, including cardiomegaly, variable fat deposition and interstitial fibrosis. SMN knockdown in human cardiomyocytes induced a metabolic shift and widespread transcriptional dysregulation, with pathway analyses identifying selective upregulation of PTEN signaling. Elevated PTEN protein levels were observed in a subset of human SMA hearts and in early postnatal hearts of *Smn^2B/−^* mice. Our results demonstrate that the heart remains a biologically relevant target of SMN deficiency and highlights cardiomyocyte-specific metabolic and PTEN signaling alterations as potential contributors to cardiac involvement in SMA.

## Introduction

Spinal Muscular Atrophy (SMA) is a genetic neuromuscular disorder marked by progressive motor neuron degeneration resulting in muscle weakness and atrophy. Mutations in the *SMN1* gene lead to low levels of the Survival Motor Neuron (SMN) protein, which is essential not only for motor neuron health but also for critical cellular functions in various tissues^1^. SMN protein supports mRNA regulation, cytoskeletal dynamics, and cellular stress response—processes vital for many cell types, including cardiac cells. While SMA’s hallmark is motor neuron degeneration, recent studies suggest that SMN deficiency may also impact cardiac health, with SMA patients showing an increased incidence of arrhythmias, conduction defects, structural cardiac abnormalities, and cardiomyopathy^2^. Supporting this concept, preclinical studies demonstrate that reduced SMN level disrupt cardiac contractility and calcium handling in model systems, indicating a mechanistic basis for cardiac dysfunction^3^. This potential cardiac involvement highlights the importance of studying cardiac pathology in SMA as part of understanding the disease’s systemic burden^4^.

In recent years, new therapies have transformed SMA patient outcomes. For instance, *nusinersen* (Spinraza), is administered via intrathecal injections and works by modifying the splicing of *SMN2* to increase SMN protein production^5^. *Onasemnogene abeparvovec* (Zolgensma), a one-time intravenous gene therapy targeted to neonates and young children, is designed to deliver a functional copy of the *SMN1* gene to increase SMN protein production throughout the body^6^. *Risdiplam* (Evrysdi), an oral SMN2 splicing modifier, increases SMN protein levels with broad distribution^7^. These therapies have dramatically improved motor function, slowed disease progression, and extended life expectancy for SMA patients, transforming SMA from a disease with high infant mortality into a manageable chronic condition for many. However, as patients live longer, they may develop complications outside of motor function, such as cardiac issues, which current therapies were not specifically designed to address. This makes it increasingly important to study cardiac health in SMA, ensuring that evolving therapeutic approaches address the full spectrum of potential central nervous system and periphery for patients with SMA.

Our study investigates heart pathology in SMA and examines how SMN deficiency specifically affects the function of human cardiomyocytes. By examining autopsy reports and cardiac tissues from patients with SMA and supplementing data from cellular and mice experiments, we explore the extent of cardiac involvement in severely affected SMA patients and identify potential biomarkers for cardiac stress related to SMN deficiency. This approach may help elucidate the contribution of SMN deficiency to cardiac function and inform future strategies to monitor the disease’s broader impact on patient health.

## Results

### Postmortem Analysis Identifies Variable Cardiac Involvement in SMA Patients

To assess cardiac abnormalities in the patients with SMA, we analyzed postmortem cardiac tissue from 14 pediatric patients with SMA type 1 who underwent comprehensive or focused clinical and/or research autopsy (**Table 1**). Consistent with advanced disease in the pre-treatment era, systemic complications were common. Clinically evident cardiovascular dysfunction including pulmonary hypertension, acute and chronic heart failure, and intermittent cardiac arrhythmia in the setting of systemic illness (bradycardia, bundle branch block and ventricular tachycardia), was present prior to death in a subset of patients, indicating that cardiac abnormalities contributed to disease burden. Among the 12 patients with full autopsies, comprehensive cardiac and systemic findings are summarized in **Tables 2, 3 and 4**. Case level review revealed substantial heterogeneity in cardiac involvement at autopsy. While most hearts appeared grossly normal, three patients exhibited increased heart mass relative to age matched expectations (i.e., cardiomegaly) and left ventricular wall thickness showed modest variability without a consistent pattern of concentric hypertrophy. In one severely affected 10 years 9 month old patient, fatty replacement of ventricular tissue was grossly evident and more significantly affected the right ventricle, in a pattern similar to that seen with arrhythmogenic right ventricular cardiomyopathy. Other grossly evident findings in a single patient each included right ventricular dilatation and prior surgical repair of congenital coarctation of the aorta (**Tables 2** and **3**). These cases variably co-occurred with pericardial effusion, pulmonary hypertension, or heart failure and spanned a range of ages and body weights, indicating that increased cardiac mass was not simply proportional to somatic growth (**Tables 2 and 4, Figure S1A**).

**Table 1.** Demographics for the cohort of 14 patients with SMA who underwent full or targeted autopsy.

| Patient ID no. | Sex | Age at death<br>(year, month) | SMN2 copies | Race | Hispanic<br>or Latino |
| --- | --- | --- | --- | --- | --- |
| 101 | M | 10y, 2m | 2 | White | No |
| 112 | F | 16y, 1m | 2 | White | No |
| 177 | F | 1y, 4m | 2 | White | No |
| 187 | F | 5y | 2 | White | No |
| 195 | M | 2y, 10m | 2 | White | No |
| 196 | F | 3y, 9m | 2 | White | No |
| 206 | M | 2y, 4m | 2 | White | No |
| 217 | M | 2y, 9m | 2 | White | No |
| 251 | M | 1y, 3m | 2 | White | No |
| 272 | M | 4m | 2 | White | Yes |
| 351 | M | 6m | 2 | White | No |
| 353 | F | 1y, 9m | 2 | White | No |
| 403 | M | 2y, 3m | 3 | African American | No |
| 617 | F | 7y, 2m | 2 | White | Yes |

**Table 2.** Patient Characteristics and Gross and Microscopic Cardiac Pathology.

| ID# | Heart weight (g) | Heart expected (mean, g) | Body Weight kg (% for age) | Cardiac Pathology | Perimortem Clinical Context |
| --- | --- | --- | --- | --- | --- |
| 101 | 162.3 | 116 | 21.4 (< 3rd%) | Pericardial effusion; cardiomegaly and myocardial fibrosis; abundant epicardial, intra-cardiac and perivascular fat. Mild but diffuse interstitial fibrosis and fibrofatty replacement of cardiac muscle. | Trach and vent dependent; acute heart failure with cardiac arrest, pneumonia, sepsis. Elective withdrawal of support. |
| 177 | 41.5 | 48 | 9.4 (3-10th%) | Evidence of prior PDA ligation and surgical correction of aortic coarctation with suture scar just distal to the third aortic branch. | Progressive respiratory failure. Elective withdrawal of support in setting of respiratory illness with acute respiratory failure. |
| 187 | 84 | 85 | Not recorded | Small pericardial effusion; Hypertrophic change of both ventricles. Mild interstitial fibrosis. Mild epicardial and intracardiac fat accumulation. | Quadriplegia; trach and vent dependent; Elective withdrawal with sepsis and acute heart failure, tachyarrhythmias, encephalopathy. |
| 195 | 61.6 | 72 | 14 (50th%) | No evident gross or microscopic abnormalities. | Progressive respiratory failure (BIPAP dependent) Withdrawal of support in setting of acute respiratory failure with viral respiratory illness |
| 196 | 65.9 | 66 | 12.5 (5-10th%) | Right ventricle with interstitial edema; Mild interstitial fibrosis and perivascular fat. | Progressive respiratory failure (BIPAP dependent) acute bronchopneumonia with cardiorespiratory arrest and resuscitation; withdrawal of support |
| 206 | 66.3 | 56 | 12.8 (50-75th%) | Mild endocardial thickening. | Progressive respiratory failure (BIPAP dependent), dysautonomia; acute respiratory failure post elective withdrawal of support |
| 217 | 106 | 57.5 | 14.2 (25-50th%) | Mild medial hypertrophy of peripheral pulmonary arteries, cardiomegaly, serous pericardial effusion. Mild interstitial fibrosis with endocardial fibrous thickening. | Progressive respiratory failure (BIPAP dependent); pulmonary HTN; withdrawal of support in setting of pneumonia with acute respiratory and cardiac failure |
| 251 | 49 | 56 | 11.8 (75-90th%) | Endocardial fibrous thickening close to valves. Mild interstitial fibrosis. Mild perivascular fat. | Progressive respiratory failure; acute respiratory failure in setting of upper respiratory illness following elective withdrawal of support |
| 351 | 35.4 | 31 | 7.55 (25th%) | None. | Chronic respiratory failure; unwitnessed death during sleep in setting of acute respiratory illness |
| 353 | 46 | 58 | Not recorded | Endocardial fibrous thickening and excess perivascular collagen deposition. Mild interstitial fibrosis. Excess fat in the conducting system and around vessels. | Chronic respiratory failure with increasing noninvasive ventilator support; elective withdrawal in setting of acute respiratory illness |
| 403 | 105 | 66.5 | 14.1 (75th%) | Cardiomegaly. Perivascular, epicardial and intracardiac fat accumulation unexpected for age. | Progressive respiratory failure (BiPAP dependent). Acute cardiorespiratory arrest with delayed resuscitation; support withdrawn |
| 617 | 106 | 142.5 | 20.9 (10-25th%) | Mild right ventricular dilatation; mild endocardial thickening involving the right ventricle; mild myxomatous degeneration mitral valve, mild interstitial and replacement fibrosis and patchy focal increase in interstitial macrophages. Increased epicardial and perivascular fat. | Trach and vent dependent; quadriplegic, pulmonary HTN, recurrent bradycardia, reduced heart rate variability, encephalopathy, support withdrawn in setting of recurrent acute respiratory failure, pneumonia/bronchitis. |
| 112 | N/A | N/A | 47.3 (5-10%) | No evident gross cardiac abnormalities. Microscopy not performed. Frozen heart samples collected rapid research autopsy | Pulmonary HTN. Full quadriplegia and prolonged respiratory support with BIPAP. Support withdrawn in setting of sepsis with heart failure, arrhythmia. |
| 272 | N/A | N/A | 4.69 (<5th%) | No evident gross cardiac abnormalities. Microscopy not performed. Frozen heart samples collected at rapid research autopsy | Progressive respiratory failure. Acute cardiorespiratory arrest at home with resuscitation. Elective withdrawal of ventilator support. |

**Table 3.** Complete systemic autopsy findings in the subset of 12 patients with SMA.

| **Systemic Autopsy Findings** | **Patients (n = 12)** |
| --- | --- |
| **Head and Neck** |  |
| Craniofacial abnormality | **10** (83%) |
| Tracheostomy | **3** (25%) |
| **Infectious & Inflammatory** |  |
| Pneumonia/chronic pneumonitis/bronchitis | **9** (75%) |
| Tracheitis | **2** (17%) |
| Esophagitis | **1** (8%) |
| Hepatic portal triaditis | **2** (17%) |
| **Edema & Fluid Accumulation** |  |
| Peripheral edema/lymphedema | **10** (83%) |
| Pleural effusion | **3** (25%) |
| Pericardial effusion | **3** (25%) |
| Ascites | **2** (17%) |
| **Coagulation & Hematologic** |  |
| Coagulopathy (+/-sepsis) | **2** (17%) |
| **Metabolic & Endocrine** |  |
| Nephrocalcinosis | **4** (33%) |
| Adrenal hypoplasia or atrophy | **4** (33%) |
| Necrosis of pancreatic endocrine cells | **1** (8%) |
| Pancreatic islet cell hyperplasia | **2** (17%) |
| **Lymphatic & Immune Dysregulation** |  |
| Lymphoid hyperplasia or malformation | **5** (42%) |
| **Cardiovascular** |  |
| Features of chronic pulmonary hypertension (HTN) | **3** (25%) |
| Supporting evidence for acute or chronic heart failure | **3** (25%) |
| **Congenital & Developmental** |  |
| Coarctation of the aorta with patent ductus arteriosus | **1** (8%) |
| Undescended testes | **3** (43% of males) |
| Ovarian follicle cysts | **2** (40% of females) |
| **Hepatic & GI Disorders** |  |
| Fatty liver, microsteatosis; hepatocyte vacuolization or cholestasis | **5** (42%) |
| Gastrotomy tube +/- fundoplication | **9** (75%) |
| Accessory spleens | **2** (17%) |
| Gallstones | **1** (8%) |
| **Musculoskeletal Abnormalities** |  |
| Skeletal muscle atrophy, fatty replacement with variable limb contractures and chest wall deformity (pectus carinatum/excavatum) | **12** (100%) |
| Thoracic scoliosis | **2** (17%) |

**Table 4.** Gross anatomical cardiac observations in 12 patients with complete autopsies.

| Anatomical features | Normal | Increased | Decreased |
| --- | --- | --- | --- |
| Heart weight | 9 (75%) | 3 (25%) | - |
| LV wall thickness | 7 (58%) | 4 (33%) | 1 (8%) |
| RV wall thickness | 12 (100%) | - | - |
| Valvular anatomy | 12 (100%) | - | - |
| Coronary arteries | 12 (100%) | - | - |
| Pericardium | 12 (100%) | - | - |

Other features included right ventricular dilatation (N=1) grossly evident fat infiltration of ventricles (N=1) and congenital cardiac structural abnormalities s/p surgical repair (N=1)

At the microscopic level, recurrent but subtle myocardial abnormalities including perivascular fat and variable interstitial fibrosis were present even in hearts classified as structurally normal (**Figure 1A-F**, **Figure S1B-E**). Among the 12 SMA hearts with complete microscopic examination, mild interstitial fibrosis was present in 7/12 (58%) and excess perivascular or intracardiac fat deposition in 7/12 (58%), with one case showing fat significantly infiltrating both cardiac ventricles (**Table 2**). It is important to note that mild and patchy perivascular and intracardiac fat can occur in children who are inactive, while our control group was made up entirely of apparently normally active infants or children prior to their deaths.

**Figure 1.**
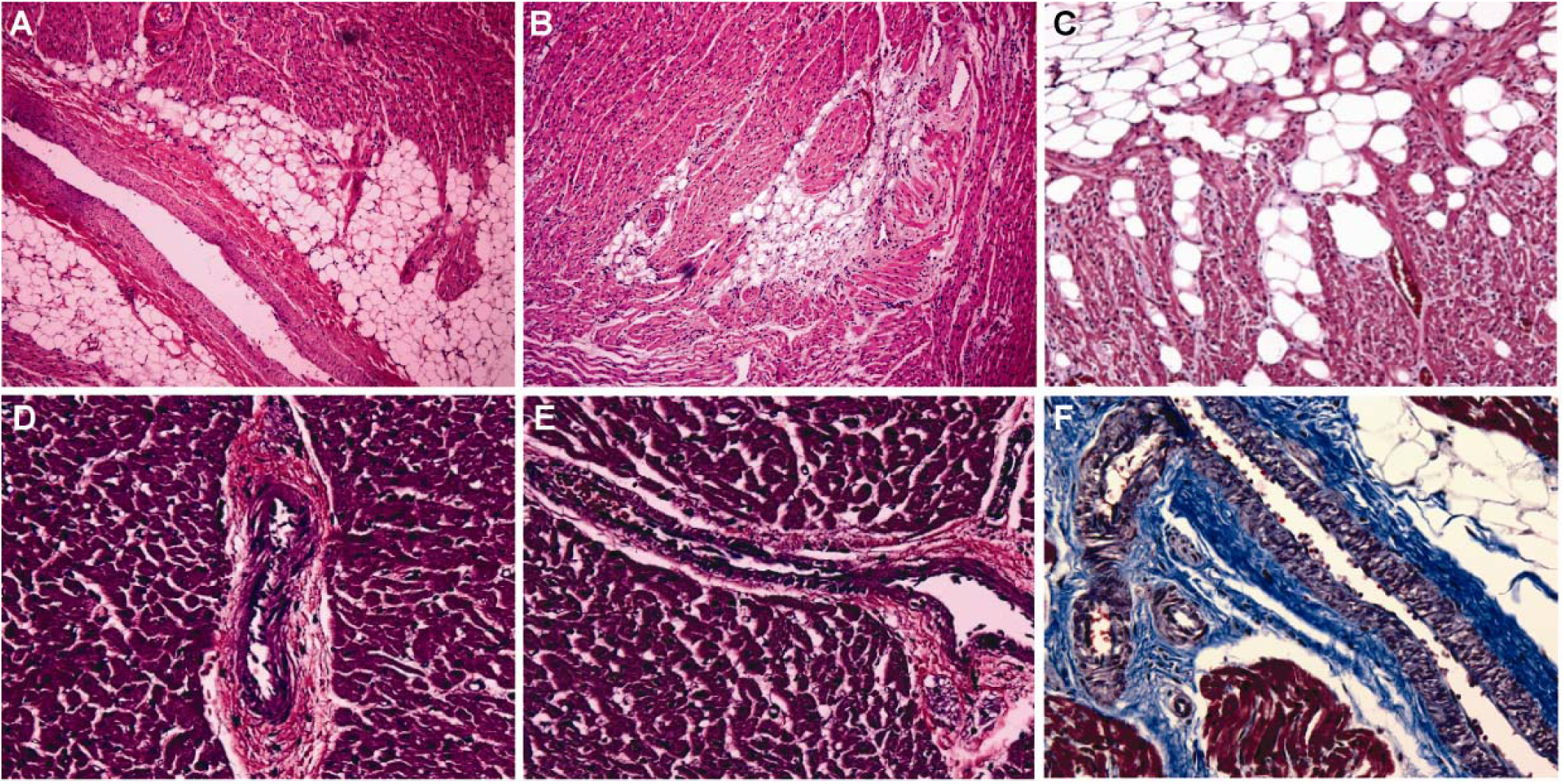
Heterogeneous cardiac fat deposition and interstitial remodeling in SMA type. **I.** A subset of patients showed variably increased perivascular fat and collagen deposition as compared to age matched controls. Pathologic findings were heterogeneous and not present in all hearts. Perivascular fat demonstrated in representative slides from **A** and **B:** 353 and 196 (shown at 10× H&E) with adipocytes surrounding arteriole and venule respectively, as compared to age matched control 4379 in **D** and **E** (40× H&E). Fat infiltrating cardiac ventricular muscle is shown in **C**: 403 (4× H&E) considered unusual given age of 2 yrs 3 months. Increased perivascular interstitial tissue is shown in **F**: 101 (40× oil trichrome), a 10 year, 9-month-old male with the most grossly evident and diffuse cardiac pathology.

Collectively, these data indicate that although overt cardiac pathology is not universal in untreated SMA type 1 patients, a subset exhibit increased cardiac mass and more subtle myocardial changes consistent with metabolic dysfunction and cardiac remodeling. These observations align with prior reports describing heterogeneous but meaningful cardiac involvement in severe, early-onset SMA and support the concept that the heart is a clinically relevant secondary organ impacted by SMN deficiency^8,9^.

### SMN Knockdown Induces Metabolic and Transcriptional Alterations in Human Cardiomyocytes

To investigate whether reduced SMN levels directly alter cardiomyocyte homeostasis, we used healthy human cardiomyocytes transduced with an AAV2 vector expressing siSMN to achieve targeted SMN knockdown (**Figure 2A**). We first assessed cellular bioenergetics using Seahorse metabolic flux assays to quantify oxygen consumption rate (OCR), extracellular acidification rate (ECAR), and estimate total ATP production (**Figure 2A**). Cardiomyocytes with decreased SMN levels exhibited a modest but consistent increase in baseline OCR accompanied by a reduction in ECAR, indicating a shift in basal metabolic activity (**Figure 2B-C**). Despite these changes, total ATP production remained unchanged following SMN depletion (**Figure 2D**). Analysis of ATP source contribution revealed increased mitochondrial ATP production with a corresponding decrease in glycolytic ATP generation, suggesting a redistribution of metabolic reliance toward oxidative phosphorylation at baseline. Although the functional consequences of this shift remain unclear, these findings are consistent with altered metabolic homeostasis and may reflect an intrinsic mitochondrial stress response that is compensated by reduced glycolytic contribution.

**Figure 2.**
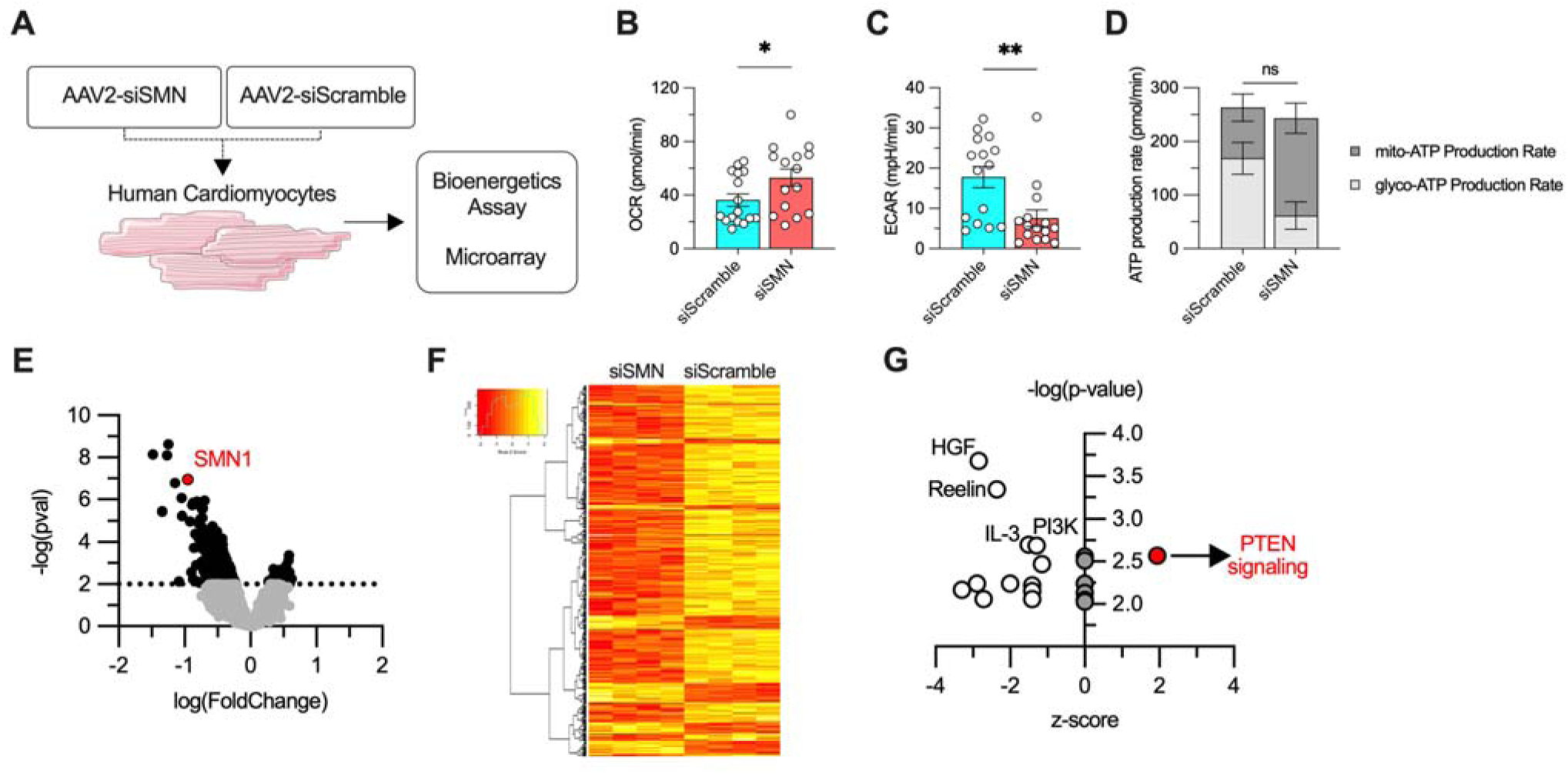
SMN knockdown alters cardiomyocyte bioenergetics and gene expression in vitro. **A:** KO schematic in cardiomyocytes **B-D:** Oxygen consumption rate (OCR) is significantly increased in siSMN-treated cardiomyocytes compared with siScramble controls and extracellular acidification rate (ECAR) is significantly decreased in siSMN-treated cardiomyocytes. n = 16 technical replicates. * p <0.05. **p < 0.01. Mitochondrial and glycolytic ATP production rates show no significant difference between siSMN and siScramble conditions (ns). **E:** Volcano plot of differential gene expression following SMN knockdown. **F:** Heatmap and hierarchical clustering of differentially expressed genes demonstrate distinct transcriptional profiles between siSMN and siScramble cardiomyocytes. **G:** Pathway enrichment analysis of differentially expressed genes identifies significant perturbation of multiple signaling pathways, including enrichment of PTEN signaling.

To further define molecular pathways affected by SMN depletion, we performed gene expression microarray analysis comparing SMN knockdown cardiomyocytes with control cells (**Figure 2A**). This analysis identified more than 1,000 differentially expressed genes, with the majority showing downregulation following SMN reduction (**Figure 2E-F** and **Table S2**). Pathway enrichment analysis revealed several significantly altered signaling networks, including HGF, Reelin, IL-3, PI3K, and PTEN signaling pathways (**Figure 2G** and **Figure S2**). Notably, PTEN signaling was the only pathway exhibiting a positive activation score, suggesting relative upregulation in response to SMN knockdown (**Figure 2G**).

To determine whether these transcriptional effects were specific to cardiomyocytes, we performed parallel SMN knockdown experiments in human myotubes as a model of skeletal muscle (**Figure S3A**). Although efficient SMN depletion was achieved, gene expression analysis revealed minimal transcriptional changes compared to controls, with no robust pathway-level alterations detected (**Figure S3** and **Table S3**). This contrast highlights differential cellular sensitivity to SMN loss and suggests that myotubes may not be particularly vulnerable to SMN-dependent metabolic and transcriptional dysregulation to the same extent observed in cardiomyocytes.

### PTEN Dysregulation in a Subset of Human SMA Hearts and Early-Stage SMA Mice

Motivated by the identification of PTEN signaling as an upregulated pathway in SMN-depleted cardiomyocytes, we next examined PTEN protein levels in available human SMA heart autopsy samples that could be fresh-frozen and analyzed by immunoblot. While PTEN expression was not uniformly elevated across all cases, a subset of SMA hearts showed clearly increased PTEN levels compared with age-matched healthy control autopsies (**Figure 3A-B**). Specifically, among seven SMA cardiac samples analyzed, three exhibited clear PTEN elevation that was not observed in controls, with the most notable increases observed in the youngest (4 months old) and oldest (16 years old) patients in this cohort (**Figure 3A-B**). These findings suggest inter-individual variability in PTEN regulation in human SMA hearts but provide direct evidence that PTEN upregulation occurs in a subset of patients.

**Figure 3.**
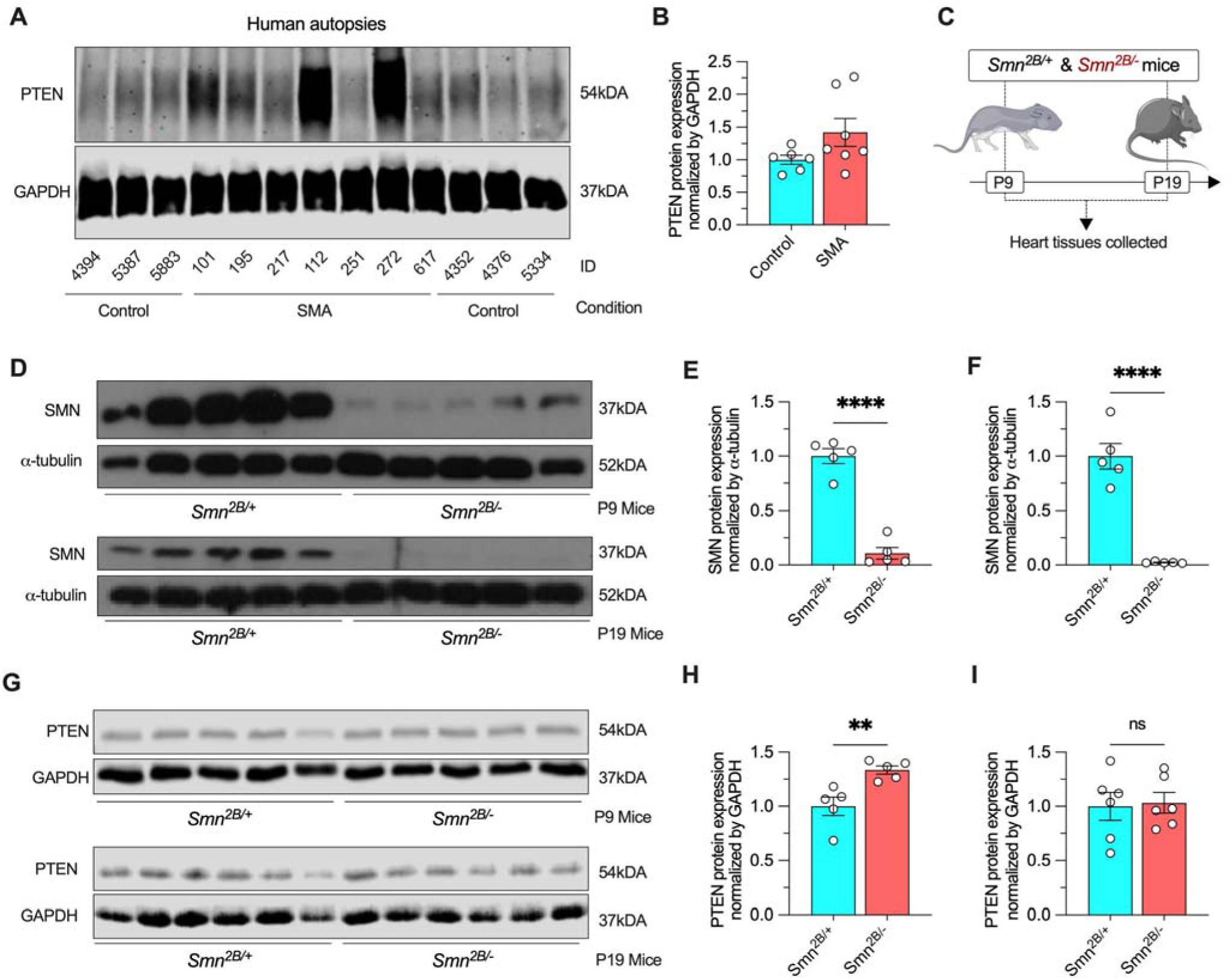
PTEN protein expression is altered in SMA human hearts and in a mouse SMA model. **A:** Immunoblots of PTEN protein expression in human cardiac tissue from control and SMA autopsy samples. **B:** Quantification of PTEN protein expression in human hearts normalized to GAPDH. n = 6-7. **C:** Schematic of the mouse SMA model (Smn^2B/+^ and Smn^2B/−^ mice) and experimental timeline, with heart tissues collected at postnatal day 9 (P9) and postnatal day 19 (P19). **D:** Immunoblots of SMN protein expression in hearts from mouse models. **E-F**: Quantification of cardiac SMN protein expression normalized to α-tubulin at (**E**) P9 and (**F**) P19. n = 5 mice. * p <0.05. **G:** Immunoblots of PTEN protein expression in hearts from Smn^2B/+^ and Smn^2B/−^ mice at P9 and P19, with GAPDH as a loading control. **H-I**: Quantification of cardiac PTEN protein expression normalized to GAPDH shows significantly increased PTEN levels in Smn^2B/−^ mice at (**H**) P9, with no significant difference at (**I**) P19. n = 5 mice.

To further assess whether PTEN dysregulation is associated with SMN deficiency in vivo, we analyzed cardiac tissue from the *Smn^2B/-^*mouse model of SMA, which exhibits systemic disease manifestations during progression (**Figure 3C**). We first confirmed reduced SMN protein levels in the hearts of *Smn^2B/-^*mice at postnatal day 9 (P9), corresponding to an early disease stage, and at postnatal day 19 (P19), when mice are more symptomatic (**Figure 3C-F**). Consistent with disease progression, SMN expression was higher at P9 and declined by P19. We then assessed PTEN protein levels in these same cardiac samples across developmental stages (**Figure 3G-I**).

PTEN levels were modestly but consistently increased in *Smn^2B/-^* hearts at the P9 stage compared with littermate controls, whereas no significant difference in PTEN expression was observed at P19 (**Figure 3G-I**). This temporal pattern suggests that PTEN upregulation is most prominent during early postnatal stages, when SMN levels are relatively higher and play a critical role in cardiac development. Over time, as SMN expression declines and developmental programs mature, PTEN levels appear to normalize. These findings support a model in which SMN deficiency is associated with stage-dependent PTEN dysregulation in the heart, with early developmental windows appearing particularly sensitive to SMN-dependent signaling perturbations.

## Discussion

In this study our multi-level dataset—spanning human postmortem hearts, SMN-deficient human cardiomyocytes, and an SMA mouse model—demonstrates that the heart remains a critical, biologically relevant target of SMN deficiency. Here, we provide the first detailed description of gross and microscopic cardiac pathology at autopsy for a cohort of type 1 patients. Our data support heterogeneous cardiac involvement in patients as well as intrinsic, stage-dependent cardiac vulnerability to SMN loss in SMA. It is important to note that in most of the postmortem hearts from the SMA patients examined, microscopic abnormalities were mild and overlap with those observed in children with prolonged inactivity related to chronic medical conditions. It is impossible to know how much the profound immobility in this severely affected early infantile onset SMA cohort impacted our observations since prior detailed microscopic cardiac pathologic examinations in genetically proven SMA type 1 patients have not been previously published. The reported clinical and gross pathologic features in our cohort align with prior clinical observations and systematic reviews reporting a range of abnormalities including congenital structural abnormalities such as septal defects and outflow-tract abnormalities and arrythmias in SMA type 0 and type 1 patients^10^, alterations in left ventricular strain in children with later onset SMA^11^, and ECG abnormalities, arrythmias and altered cardiac function in later onset SMA patients.^2,12^

Cardiomyocyte SMN knockdown produced a compensatory metabolic shift rather than ATP failure, characterized by increased baseline oxidative phosphorylation and reduced glycolytic contribution without altering total ATP production. These findings complement recent work in SMA mouse models showing disrupted cardiac fatty acid metabolism and altered mitochondrial function associated with functional cardiac impairment^3^ as well as reports of electrical changes (e.g., bradycardia, deep Q waves) in patients, supporting a spectrum from subclinical remodeling to clinically apparent disease. PTEN activation emerged as a shared SMN-sensitive pathway and was elevated in early postnatal hearts, consistent with developmental sensitivity and with reports that PTEN suppression mitigates SMA severity in mice^13,14^. Importantly, our data extend these reports by coupling human autopsy findings with mechanistic analyses, supporting the concept that cardiac involvement in SMA also reflects intrinsic tissue vulnerability rather than solely secondary effects of neuromuscular failure^15^, which are obviously also present.

This pattern suggests a compensatory redistribution of energy metabolism rather than overt bioenergetic failure, consistent with an early or subclinical stress response. Together with prior reports of impaired calcium handling and SERCA2 dysregulation in SMA cardiomyocytes derived from mice and patient iPSCs, our results reinforce the notion that SMN deficiency affects multiple interconnected pathways that are essential for cardiomyocyte function.

A key finding of our transcriptomic analyses was the identification of PTEN signaling as the main pathway with predicted activation following SMN knockdown in human cardiomyocytes. This observation is notable in light of prior studies implicating PTEN as a disease modifier in SMA. Godena and Ning (2017) proposed PTEN as a therapeutic target in SMA based on its role in regulating neuronal survival and growth, and Little et al. (2015) demonstrated that PTEN depletion ameliorates disease severity and modestly prolongs survival in SMA mouse models^13,14^. Our data extend this framework to the heart, showing that PTEN upregulation occurs in a subset of human SMA cardiac tissues and is recapitulated in early postnatal stages in the *Smn^2B/−^* mouse heart. These findings suggest that PTEN dysregulation may represent a shared SMN-sensitive pathway across tissues, while also highlighting important temporal and tissue-specific differences in PTEN regulation.

The stage-dependent nature of PTEN upregulation observed in SMA mouse hearts is particularly informative. PTEN elevation was evident at early postnatal stages (P9), when SMN expression is relatively high and cardiac developmental programs are still active, but not at later symptomatic stages (P19), when SMN levels decline and PTEN expression normalizes. This temporal pattern suggests that SMN deficiency may disrupt developmental signaling thresholds rather than driving sustained PTEN activation throughout disease progression. Such a model is consistent with the heterogeneous PTEN expression observed in human SMA hearts and may help reconcile why PTEN modulation is beneficial in some contexts but not universally altered across all tissues or disease stages. Importantly, our findings also indicate that cardiomyocytes exhibit marked transcriptional and metabolic sensitivity to SMN loss, underscoring cell-type–specific dependencies on SMN-regulated pathways, with PTEN-linked signaling contributing early and heterogeneously.

Finally, these results have important implications for SMA patients in the era of disease-modifying therapies. While treatments such as nusinersen, onasemnogene abeparvovec, and risdiplam have dramatically improved motor outcomes and survival^5–7^, they were not specifically designed to address cardiac or other systemic manifestations of SMN deficiency. In the treatment era, routes of SMN restoration differ in biodistribution. It remains to be seen whether the broad tissue distribution of risdiplam (an orally administered RNA modifying therapy increasingly used across age cohorts from prenatal to adults) and intravenously administered onasemnogene abeparvovec (zolgensma) (with presumed variable systemic transduction in neonates and young children), provides superior multi-organ protection compared to localized CNS-targeting therapies, or how it may impact PTEN or other potentially relevant biomarker levels. The newly approved intrathecal formulation of onasemnogene abeparvovec (Ivitsma), while attractive in reducing potential immune-mediated liver toxicity, could potentially leave cardiomyocytes and other peripheral tissues more vulnerable to SMN deficiency in the setting of decreased systemic exposure. As individuals with SMA live longer, how these differing treatment modalities influence long-term cardiac biology is unknown, and are dependent on their SMN2 copy number, age of administration and systemic vs targeted CNS exposure. Our data suggest that cardiomyocyte metabolic stress and PTEN-dependent signaling alterations may represent early indicators of cardiac involvement and potential targets for adjunctive therapeutic strategies. Future studies will be required to determine whether cardiac phenotypes persist or evolve in treated SMA populations and whether early modulation of metabolic or PTEN-related pathways can mitigate long-term cardiac risk.

How differences in tissue exposure (intrathecal vs. systemic) translate into cardiac protection or biomarker modulation (e.g., PTEN-related signatures) is not yet known; emerging reviews on treated populations and combination strategies underscore uncertainty about long-term extra-CNS effects and the need for standardized outcomes beyond motor function. Our data suggest that cardiomyocyte metabolic stress and PTEN-linked signaling may be early indicators of cardiac involvement in SMA and candidate targets for adjunctive strategies, particularly for individuals with severe genotypes (≤2 SMN2 copies) who survive longer with therapy. Prior PTEN-modulation studies in SMA provide proof-of-concept for pathway tractability; however, any attempt to modulate PTEN in humans must balance potential benefit with PTEN’s role as a tumor suppressor and its broad biology. Prospective studies in contemporary, treated SMA cohorts should (i) incorporate cardiac phenotyping (ECG, echocardiography, myocardial strain, biomarkers and, where feasible, cardiac MRI), (ii) evaluate circulating or imaging biomarkers of metabolic stress and PTEN-pathway activity, and (iii) define whether early pathway modulation mitigates long-term cardiac risk as survival improves.

## Methods

### Ethics, Subjects, and Study Design

Sex was not considered as a biological variable in this study. Ethical approval and written informed parental consent were obtained for all human participants via Institutional Ethics Review Board approved research protocols at the University of Utah (UU IRB_00008751 from 2008-2015 and the Massachusetts General Hospital (MGH IRBs 2015P001934 and 2016P000469 from 2016-2022). Autopsies were performed in collaboration with pathologists at Primary Children’s Medical Center, Salt Lake City, UT, USA; and Massachusetts General Hospital, Boston, MA, USA. All procedures adhered to the ethical principles and guidelines of the WHO Guiding Principles on Human Cell, Tissue and Organ Transplantation.

Mouse studies were approved by the Animal Care and Veterinary Services of the University of Ottawa, Ontario, Canada (protocols #OHRI-1927 and #OHRI-1948) and were conducted in accordance with institutional and national standards for the care and use of laboratory animals.

### Animals

*Smn^+/−^* mice were bred with *Smn^2B/2B^*mice to obtain *Smn^2B/+^* and *Smn^2B/-^* progeny maintained on the C57BL6/J background^16,17^. The *Smn^2B/-^* mice are a model of severe SMA and the asymptomatic heterozygous *Smn^2B/+^* mice are used as controls in these experiments. Both male and female mice were used in the analysis.

### Human heart autopsies

This cohort study included 14 pediatric patients with genetically confirmed SMA with a severe infantile onset SMA type 1 clinical phenotype. Clinical characteristics and autopsy findings for these cases are summarized in Tables 1-3, and Table S1^18–20^. None received the FDA-approved disease-modifying therapies nusinersen, risdiplam, or onasemnogene abeparvovec. Comprehensive pre-mortem clinical data from medical records were available for all 14 patients with SMA. Initial pathologic examinations were performed following an autopsy protocol that included partial or full cardiac examination by a clinical academic pathologist at the corresponding institution. An academic cardiac pathologist at MGH reviewed additional H&E and trichrome stained slides to confirm gross microscopic features and assess the presence or absence of interstitial fibrosis and/or endocardial fibrosis. Age and sex matched control tissues were obtained from the NIH Neurobiobank at the University of Maryland. After gross inspection and dissection, tissues were collected as 1 cm³ blocks from right and left ventricles and immersion fixed in 4% paraformaldehyde or 2.5% glutaraldehyde for electron microscopy studies. In 7 of 14 SMA cases, additional cardiac tissue samples were obtained and flash-frozen for later analysis.

Cardiac tissues from patients with SMA and controls were fixed, embedded, sectioned and stained with hematoxylin and eosin (H&E) and Masson’s Trichrome. Additional slides were generated for each SMA and control sample and imaged on an Olympus BX53 microscope using 2×, 4×, 10×, 20×, 40×, and 100× oil-immersion lenses (Olympus Corporation, Shinjuko, Tokyo, Japan). Images were collected using Q Capture Pro Imaging software (Surrey British Columbia, Canada). Images were formatted using Adobe Photoshop and Microsoft Office software products. The entire slide was sampled at each magnification (4×, 10×). At the highest magnifications (40×, 100× oil), only one or two images per region of interest were collected. Sufficient frozen tissue was available in 7 SMA and 6 healthy controls for protein extraction and analysis.

### Cell culture

Cell culture experiments were performed in differentiated human cardiomyocytes and myotubes. Human cardiomyocytes (Axol Bioscience Limited, Cambridgeshire, England; Cat. No. ax2520; Lot No. 2520310317) were cultured and differentiated in coated plates using supplemented media from the Human iPSC-Derived Ventricular Cardiomyocyte Kit (Axol Bioscience Limited, Cambridgeshire, England). Primary human skeletal myoblasts (Thermo Fisher Scientific; A11440) were cultured with SkGM™-2 Skeletal Muscle Cell Growth Medium-2 Bullet Kit (Lonza, CC-3245) supplemented with 10% fetal bovine serum (FBS) and 1% pen/strep. Myoblasts were differentiated into myotubes over four days by replacing media with Dulbecco’s modified Eagle’s medium (DMEM; Gibco) supplemented with 2% horse serum and 1% pen/strep. After five days of differentiation, myotubes were maintained in SkGM™-2 Skeletal Muscle Cell Growth Medium-2 Bullet Kit (Lonza, CC-3245) supplemented with 10% FBS or patient plasma and 1% pen/strep.

To knockdown SMN in cardiomyocytes, cells were transduced with adenoviruses (MOI 100) expressing GFP and specific siRNA for a human SMN sequence (AAV2-GFP-U6-h-SMN1-shRNA) or a scramble sequence (AAV2-GFP-U6-scrmb-shRNA). All vectors were produced by Vector Biolabs (Malvern, PA). Twelve hours after transduction, the medium was replaced with fresh differentiation medium. All cells used in this study were maintained at 37°C in 5% CO2 and tested negative for mycoplasma. To knockdown SMN in myotubes, specific siRNA sequences (Thermo Fisher Scientific) were used for human SMN using Lipofectamine RNAiMAX Transfection Reagent (Thermo Fisher Scientific; 13778-075) in Opti-MEM Media (Thermo Fisher Scientific; 31985062). Transfections with scramble siRNA (4390843) were used for the control groups.

### Seahorse Assay

Oxygen consumption rate (OCR) and extracellular acidification rate (ECAR) were measured with extracellular flux analysis (XF96, Agilent Seahorse, MA, USA) in sodium bicarbonate-free DMEM supplemented with 31.7 mM NaCl, 10 mM glucose and 2 mM glutamax (pH 7.4 adjusted using NaOH) as previously described by our group^21^. Total protein content was measured with a BCA assay (Thermo Fisher Scientific, 23225) to confirm comparable protein levels at the endpoint of the Seahorse assay.

### Microarray and Pathway Analysis

Transcriptomes from human cardiomyocytes and myotubes were determined using Clariom S Assay, human (Thermo Fisher Scientific) using the Thermo Fisher Scientific facility services (Santa Clara, CA, USA) as previously described by our group^21^. All data analysis was done in R^22^ using the packages “Limma”^23^ and “Oligo”^24^ through Bioconductor. Affymetrix data were first normalized by using robust multi-array averages. These normalized data were then fit through a linear model. The empirical Bayes statistics for differential expression was used to calculate all statistical values. Graphs were generated using ggplot2^25^. Microarray data will be submitted to the NCBI Gene Expression Omnibus (GEO) and will be made publicly available upon acceptance of this manuscript

### Immunoblotting

SMN, PTEN and GAPDH protein levels were determined by immunoblotting as previously described^26^ with few modifications. Tissues were lysed and 20 μg of isolated total protein was loaded in a 4–20% precast protein gel (Biorad, #4561096) and subjected to electrophoresis. Proteins were transferred to a PVDF membrane and blocked for 1 h at room temperature in Odyssey blocking buffer (Li-Cor, Lincoln, NE). Membranes were incubated with primary antibodies overnight at 4°C. Primary antibodies were used to probe for SMN (BD Biosciences; 610647), PTEN (Cell Signaling; #9552) and GAPDH (Cell Signaling; #2118). Membranes were imaged using a ChemiDoc Touch System (Bio-Rad, USA) or the LI-COR Odyssey Infrared Imaging System (LI-COR, Inc., USA). SMN and PTEN expression were normalized to GAPDH levels.

### Statistical Analysis and Data Availability

Data are presented as mean ± standard error of the mean with dots as individual values. Sample size is indicated in the figure legends. Statistical analyses were performed using GraphPad Prism 10 software (GraphPad Software, Inc). Unpaired 2-tailed Student *t* tests were used to compare groups. Statistical significance was defined as *p* < 0.05. All data relevant to this study are contained within the article.

## Author Contributions

K.J.S. and C.R.R.A. directed the research project. R.G., L.L.H, R.G.S., E.J.E., S.D.S.D.L., A.J.J., F.C.N., J.R.S., K.J.S. and C.R.R.A collected human samples or clinical data. L.C. and R.K. performed mouse experiments. R.G., R.G.S., and C.R.R.A performed cell culture and additional experiments. R.G., L.L.H, R.G.S., E.J.E., L.C, J.R.S., R.K., K.J.S. and C.R.R.A analyzed the data. All authors participated in the data interpretation. L.L.H. and C.R.R.A. drafted the manuscript. All authors reviewed and approved the final manuscript.

## Acknowledgements

C.R.R.A received a fellowship from the MGH ECOR, a Charles A. King Trust Postdoctoral Research Fellowship, Bank of America, N.A., Co-Trustees, a James L. and Elisabeth C. Gamble Endowed Fund for Neuroscience Research / Mass General Neuroscience Transformative Scholar Award, a MGH Physician/Scientist Development Award, and a National Institutes of Health (NIH) grant K01NS134784. L.C. received a Vanier Canada Graduate Scholarship from the Canadian Institutes of Health Research. K.J.S was funded by a NIH grants R01HD054599 and R21NS108015, Biogen and Cure SMA. We are grateful to Chrystalle Katte Carreon, MD The Stella & Richard Van Praagh Cardiac Registry, Boston Children’s Hospital, Boston, MA for her help in ensuring accuracy and providing context for human cardiopathologic descriptions in figures 1 and S1. We are grateful to all the patients and families who participated in this study.

## Supplementary Material

**Table S1.** Control Subjects Characteristics

**Table S2**. List of genes from microarray performed after SMN knockdown in human **cardiomyocytes** [Excel file]

**Table S3**. List of genes from microarray performed after SMN knockdown in human **myotubes** [Excel file]

**Figure S1.**
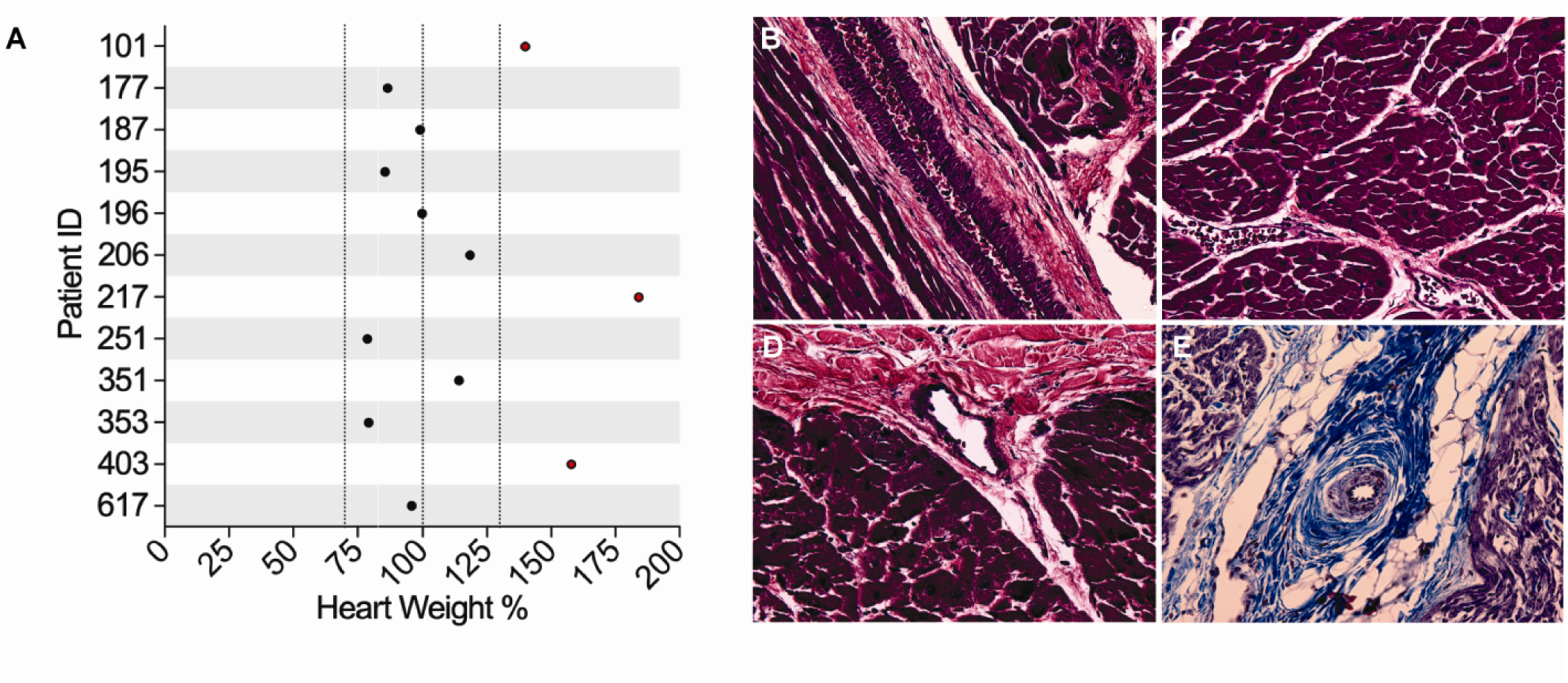
Cardiac mass variability and representative histopathologic features in SMA patients. **A:** Heart weight expressed as percent of age- and sex-expected values for individual SMA patients. The solid vertical line at 100% denotes the expected heart weight for age and sex, and the dashed vertical lines indicate ±1 standard deviation from the expected value. **B-D**: Additional histologic sections of human tissue from 12 yo male control, 40x H&E, and **E:** 40× trichrome image from 101, 10 years 9 months old male SMA patient, representative of the increased perivascular collagen and fat diffusely evident in both ventricles.

**Figure S2.**
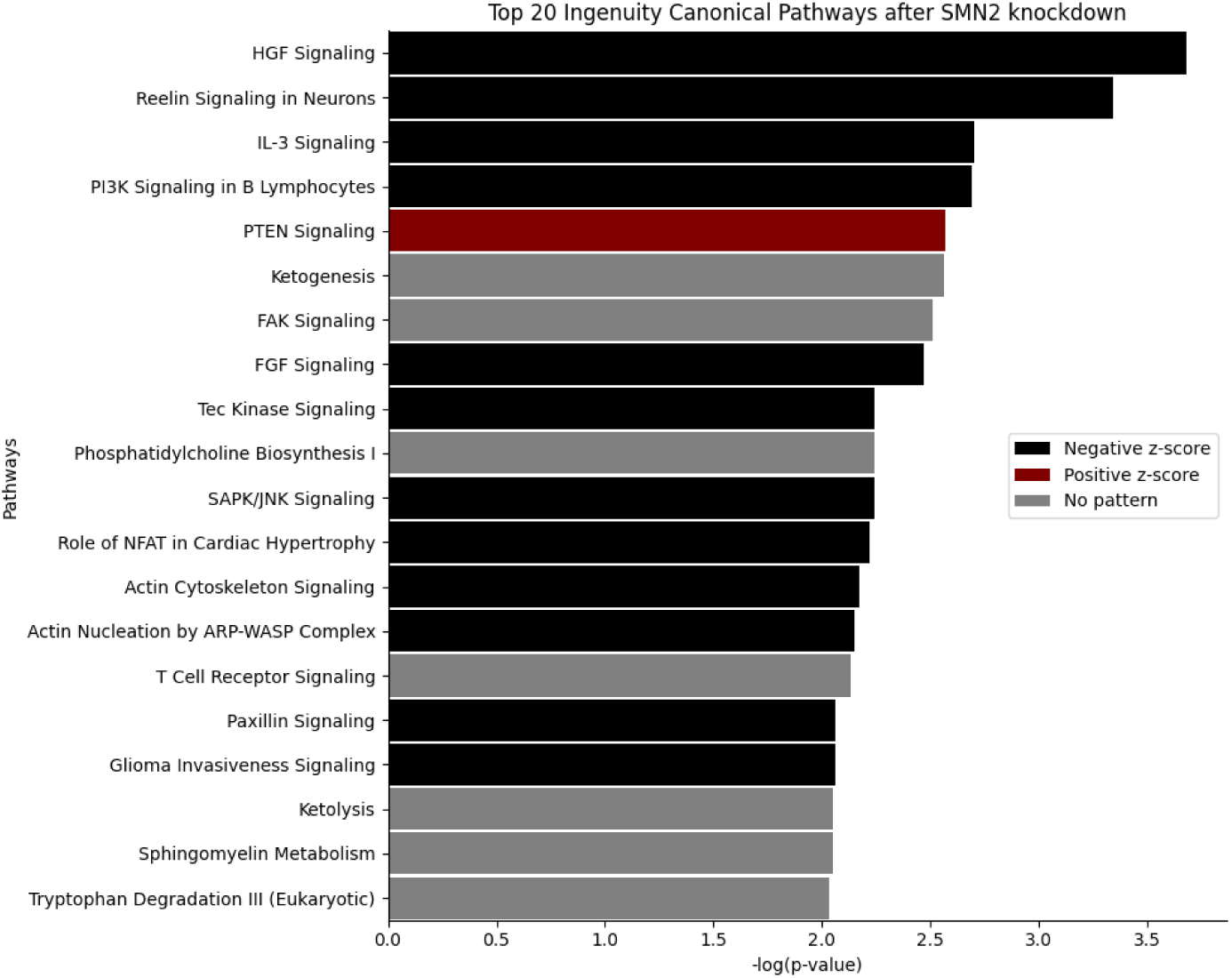
Expanded pathway analysis following SMN2 knockdown in human cardiomyocytes. Ingenuity pathway analysis of differentially expressed genes after SMN2 knockdown in human cardiomyocytes, showing the top 20 significantly enriched canonical pathways ranked by –log(p value). Bar color denotes predicted directionality based on z-score: black indicates negative z-score (predicted pathway inhibition), red indicates positive z-score (predicted pathway activation), and gray indicates no consistent activation pattern.

**Figure S3.**
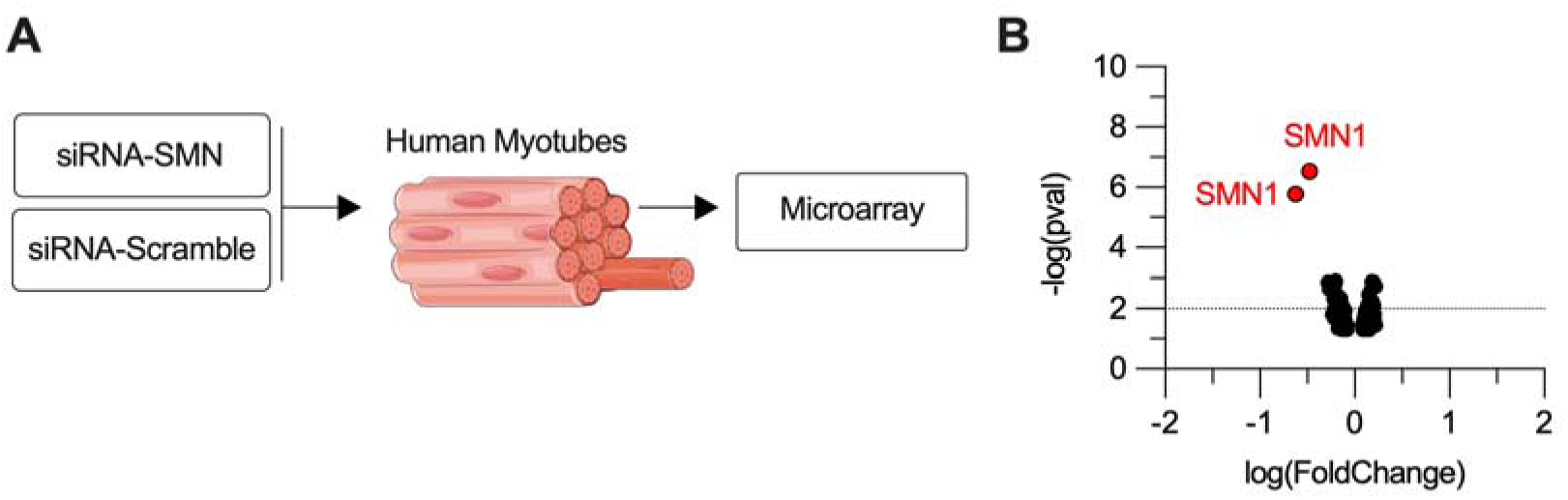
Myotubes data following SMN knockdown. **A:** Experimental schematic showing siRNA-mediated knockdown of SMN (siRNA-SMN) or scrambled control (siRNA-Scramble) in human myotubes, followed by transcriptomic analysis using microarray. **B:** Volcano plot of microarray gene expression data comparing siRNA-SMN versus control myotubes.

**Table S1.** Demographic, clinical, and autopsy characteristics of the control subjects.*

| Control ID | Age at death<br>(year, month) | Sex | Ethnicity | SMN1 | SMN2 | Cause of death | Heart<br>weight (g) |
| --- | --- | --- | --- | --- | --- | --- | --- |
| 1284 | 3y 4m | F |  | 2 | NT | Drowning |  |
| 1453 | 1y 3m | F | African American | 2 | 2 | Choking/Asthma |  |
| 1798 | 1y 4m | F | Caucasian | 2 | 1 | Bowel intussusception |  |
| 4379 | 3 y | M | Caucasian | 3 | 2 | Drowning |  |
| 1864 | 2y 6m | F |  | 2 | NT | Laryngitis and<br>Bronchiolitis |  |
| 5282 | 2y 10m | M | Hispanic | 2 | 2 | Asphyxia, choking on<br>food bolus (hot dog) | 77 |
| 5564 | 2y | M | African American | 3 | 0 | Asphyxia, choking on<br>food bolus | 110 |
| 5883 | 2 m | M | African American | 4 | 0 | Sudden infant death<br>syndrome | 40 |
| 4352 | 4 m | M |  | 2 | NT | Acute and Chronic<br>Tracheo-Bronchiolitis | 49.5 |
| 4376 | 7 m | M | Caucasian | 2 | 2 | Probable asphyxiation | 53 |
| 4394 | 3y 2m | F |  | 2 | 1 | Fever, undetermined | 65.8 |
| 5334 | 12y | M | Caucasian | 2 | 1 | Drowning | 135 |
| 5387 | 12y | M |  | 2 | NT | Drowning |  |

Adapted from *Impaired kidney structure and function in spinal muscular atrophy*, Nery, F.C. et al., Neurol. Genet. 5, e353 (2019). \*Control samples from the NIH Neurobiobank at the University of Maryland.

